# ChIPdig: a comprehensive user-friendly tool for mining multi-sample ChIP-seq data

**DOI:** 10.1101/220079

**Authors:** Ruben Esse, Alla Grishok

## Abstract

**Background:** In recent years, epigenetic research has enjoyed explosive growth as high-throughput sequencing technologies become more accessible and affordable. However, this advancement has not been matched with similar progress in data analysis capabilities from the perspective of experimental biologists not versed in bioinformatic languages. For instance, chromatin immunoprecipitation followed by next-generation sequencing (ChIP-seq) is at present widely used to identify genomic loci of transcription factor binding and histone modifications. Basic ChIP-seq data analysis, including read mapping and peak calling, can be accomplished through several well-established tools, but more sophisticated analyzes aimed at comparing data derived from different conditions or experimental designs constitute a significant bottleneck. We reason that the implementation of a single comprehensive ChIP-seq analysis pipeline could be beneficial for many experimental (wet lab) researchers who would like to generate genomic data.

**Results:** Here we present ChIPdig, a stand-alone application with adjustable parameters designed to allow researchers to perform several analyzes, namely read mapping to a reference genome, peak calling, annotation of regions based on reference coordinates (e.g. transcription start and termination sites, exons, introns, 5′ UTRs and 3′ UTRs), and generation of heatmaps and metaplots for visualizing
coverage. Importantly, ChIPdig accepts multiple ChIP-seq datasets as input, allowing genome-wide differential enrichment analysis in regions of interest to be performed. ChIPdig is written in R and enables access to several existing and highly utilized packages through a simple user interface powered by the Shiny package. Here, we illustrate the utility and user-friendly features of ChIPdig by analyzing H3K36me3 and H3K4me3 ChIP-seq profiles generated by the modENCODE project as an example.

**Conclusions:** ChIPdig offers a comprehensive and user-friendly pipeline for analysis of multiple sets of ChIP-seq data by both experimental and computational researchers. It is open source and available at https://github.com/rmesse/ChIPdig.

## Background

Interactions between nuclear proteins and DNA are vital for cell and organism function. They control DNA replication and repair, safeguard genome stability, and regulate chromosome segregation and gene expression. Chromatin immunoprecipitation coupled with high-throughput sequencing (ChIP-seq) is a powerful method for assessing such interactions and has been widely used in recent years to map the location of post-translationally modified histones, transcription factors, chromatin modifiers and other non-histone DNA-associated proteins in a genome-wide manner. This progress has been fostered by the increasing technical feasibility and affordability of this technology, with more documentation and technical support available to researchers, including commercial library preparation kits, aided by the plummeting costs of sequencing and the ease of multiplexing samples. In light of this progress, ChIP-seq data sets are continuously deposited in publicly-accessible databases, such as the National Center for Biotechnology Information’s (NCBI) Gene Expression Omnibus (GEO) and the ENCODE consortium portal [1]. Therefore, there is an unprecedented wealth of epigenomic data in the public domain that can be used for integrative and correlative analyses.

This important advancement had not been accompanied with similar progress in user-friendly post-sequencing data analysis pipelines, which is still a significant bottleneck often handled by skilled bioinformaticians. A key step in ChIP-seq data analysis is to map reads to a reference genome assembly. Programs such as BWA [2] or Bowtie/Bowtie2 [3,4] are frequently employed for this purpose. Aligned data are then processed to find regions of enrichment along the genome, thereby identifying potential loci of DNA binding by the target protein or of deposition of the histone post-translational modifications of interest. This process is known as peak calling and can be performed by using algorithms such as MACS/MACS2 [5,6] and SICER [7]. SAMtools is another popular software used in next-generation sequencing (NSG) analysis that provides various utilities for manipulating alignments, including sorting, merging, indexing and generating alignments in a per-position format [8].

Following the initial and common steps in ChIP-seq data analysis mentioned above, downstream analysis is often more customizable and may present a hurdle, especially in the case of comparing data derived from different conditions or experimental settings. Artifacts arising from bias of DNA fragmentation, variation of immunoprecipitation efficiency, as well as polymerase chain reaction (PCR) amplification and sequencing depth bias, result in ChIP-seq experiments with distinct signal-to-noise ratios and impose great challenges to the computational analysis [9]. Several methods addressing the differential enrichment analysis problem, i.e. the detection of genomic regions with changes in ChIP-seq profiles between two distinct samples or sets of replicate samples, have been proposed [10–12]. Generally, these methods rely on the initial detection of candidate peak regions by a conventional peak calling algorithm, and then this peak-defining information is applied to analysis with methods tailored for the differential expression analysis of RNA-seq data such as edgeR [13] or DESeq [14]. Downstream analysis of ChIP-seq data may also involve the annotation of genomic regions based on reference coordinates (e.g. distance from nearest transcription start site) and visualization of normalized coverage by means of heatmaps and metaplots.

Importantly, ChIP-seq data analysis typically relies, at least in part, on tools that have been designed to run primarily on Linux/Unix-based systems, while biologists who need to work with NGS may be unfamiliar with such operating systems. In the past few years, several packages designed to handle NGS data have been released for R, a popular platform-independent programming language and software environment for computing and graphics. Here we present ChIPdig, an open-source application that leverages on several R packages to enable comprehensive and modular analysis of multi-sample ChIP-seq data sets through a user-friendly graphical user interface (GUI), allowing any experimental biologist with minimal computational expertise to use it easily.

## Implementation

ChIPdig is developed in R using package Shiny and relies on multiple R/Bioconductor packages, including QuasR and BSgenome [15], for mapping reads to a reference genome assembly [16], BayesPeak for peak calling [17], csaw [12] and edgeR [13] for differential enrichment analysis, ChIPseeker for annotation of genomic regions of interest [18], and EnrichedHeatmap for generating coverage heatmaps [19]. Packages used in several analysis modules implemented in ChIPdig are GenomicRanges [20], valr [21], shinyFiles, GenomicFeatures, ggplot2, ggsignif, reshape2 and circlize. A few command lines provided in https://github.com/rmesse/ChIPdig suffice to launch the GUI from within R Studio, allowing integrative and interactive usage of these powerful libraries without requiring programming or statistical experience from the user.

## Results

ChIPdig has the following capabilities: (1) alignment of reads to a reference genome; (2) normalization and comparison of distinct ChIP-seq data sets; (3) annotation of genomic regions; (4) generation of comparative heatmaps and metaplots for visualization of normalized coverage in user-defined specific regions. Each capability corresponds to a specific analysis module which can be loaded separate from other modules and using different input files. To illustrate each module, raw single-end ChIP-seq data for H3K4me3 and H3K36me3 in the model organism *Caenorhabditis elegans* were downloaded from NCBI GEO with series IDs GSE28770 and GSE28776, respectively. These two histone modification marks have different distribution profiles, namely promoter-proximal enrichment with punctuated peaks for H3K4me3, and gene body enrichment with broad peaks for H3K36me3 [22]. Figure 1 shows the GUI displayed upon launching ChIPdig from R Studio. The left pane displays four radio buttons, each corresponding to one of the four analysis modules, as well a clickable box for selection of the folder containing the input files for the analysis.

**Fig. 1.**
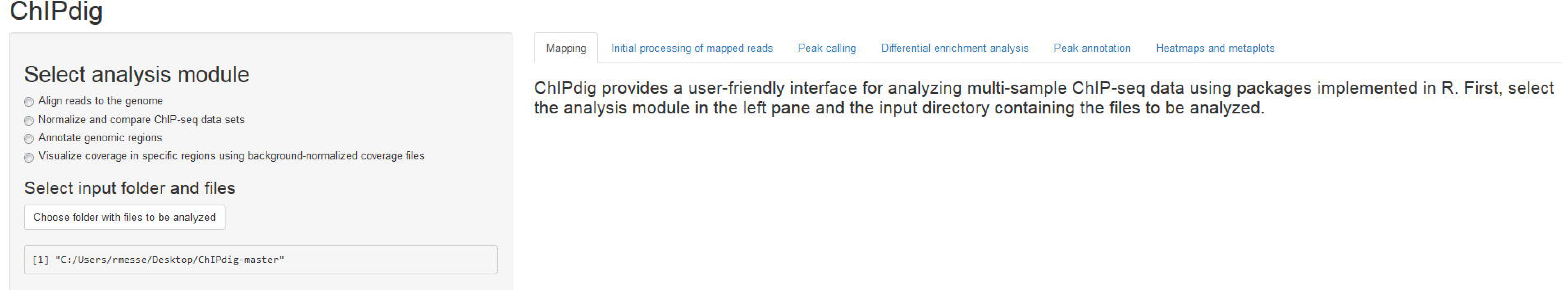
ChIPdig user interface displayed upon launch. The left pane displays four radio buttons for selection of the analysis module and a clickable box for selection of the folder containing the input files. Additional features are sequentially unlocked as the user progresses through the software suite. Outputs generated throughout the analysis and additional instructions are placed in the main panel under a tab corresponding to the analysis module selected by the user.

### • Alignment of ChIP-seq reads

The aligning module of ChIPdig relies primarily on the QuasR package, which supports the analysis of single-read and paired-end deep sequencing experiments [16]. If desired, read preprocessing can be performed to prepare the input sequence files prior to alignment, e.g. removal of sequence segments corresponding to adapters and low quality reads. Sequence files in FASTQ format for each input DNA and ChIP sample corresponding to the H3K4me3 and H3K36me3 ChIP-seq experiments were aligned to the WS220/ce10 reference genome (available through the BSgenome package [15]) (Fig. 2), producing an average of 10 million mapped reads per sample. A report with several quality metrics for each alignment was automatically generated and is available in Additional file 1.

**Fig. 2.**
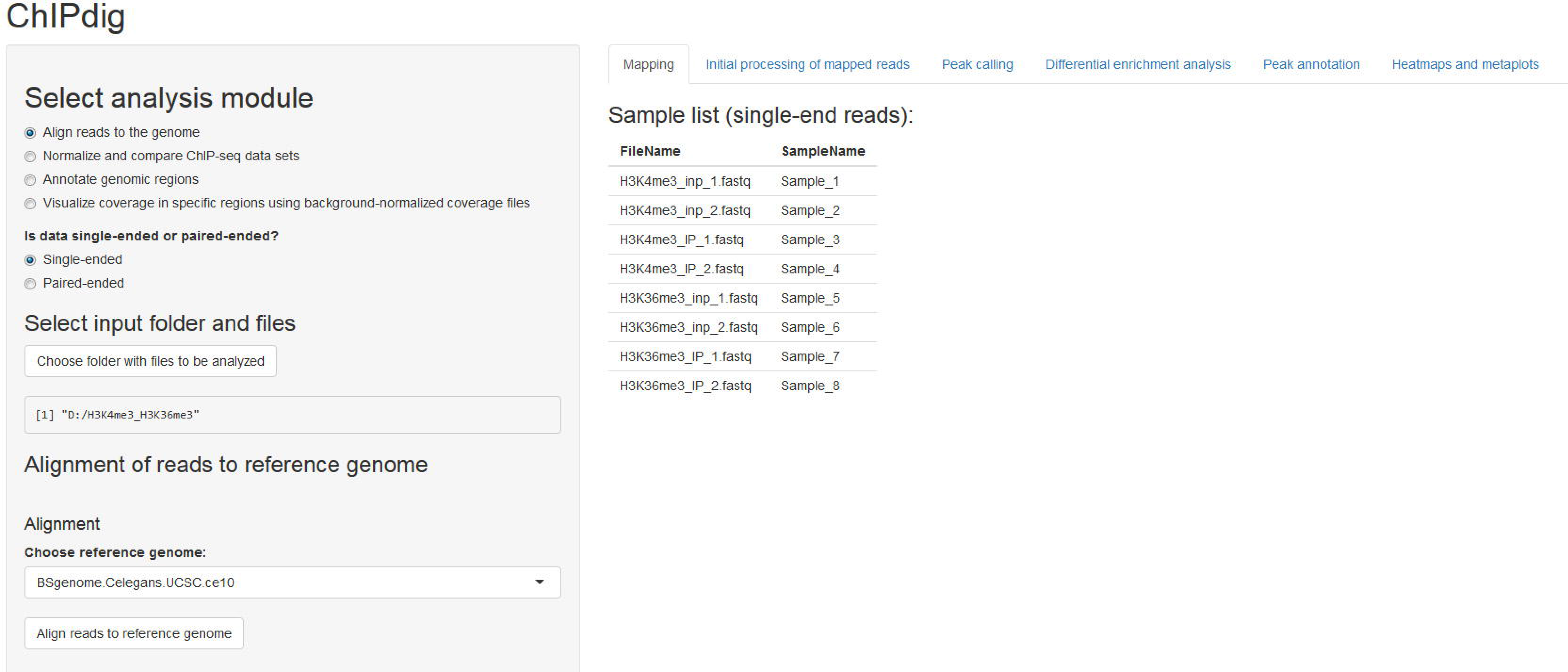
Alignment module. The user supplies a text file listing the input files corresponding to unmapped reads in either uncompressed (e.g.: ‘.fq’, ‘.fastq’) or compressed (e.g.: '‘.gz’, ‘.bz2’, ‘.xz’) format. Both single-end and paired-end data are supported. A scroll-down menu displays the reference genome assemblies available for alignment.

### • ChIP-seq data normalization, peak calling and differential enrichment analysis

A recurrent problem in ChIP-seq data analysis lies at the comparison of multiple coverage profiles generated from different experiments or corresponding to different conditions. This problem is addressed in the second analysis module of ChIPdig. Upon its selection, the user is prompted to provide a tab-delimited text file with the names of each ChIP sample (treatment) mapped reads file in BAM format, the matching input DNA (control) file, a sample ID, the condition or target name, and a color designation for the output peak and coverage files (Fig. 3a). Any of the 657 R built-in colors can be chosen (Additional file 2). A bin size parameter set to 50 bp (base pairs) by default is used in both peak calling and genome-wide library normalization for differential enrichment analysis. If desired, duplicate reads can be removed prior to normalization and sequences can be extended to a median fragment length that can be estimated either computationally or experimentally (e.g. average size of fragments generated by chromatin shearing) (Fig. 3b). The initial mapped read processing outputs a table with library sizes and median fragment sizes, a chromosome size list, and a multidimensional scaling plot representing the similarity of samples in the data set (Fig. 3c). Data are normalized based on library sizes and on the trimmed means of M values (TMM) approach [23], which is implemented in the edgeR package [13].

**Fig. 3.**
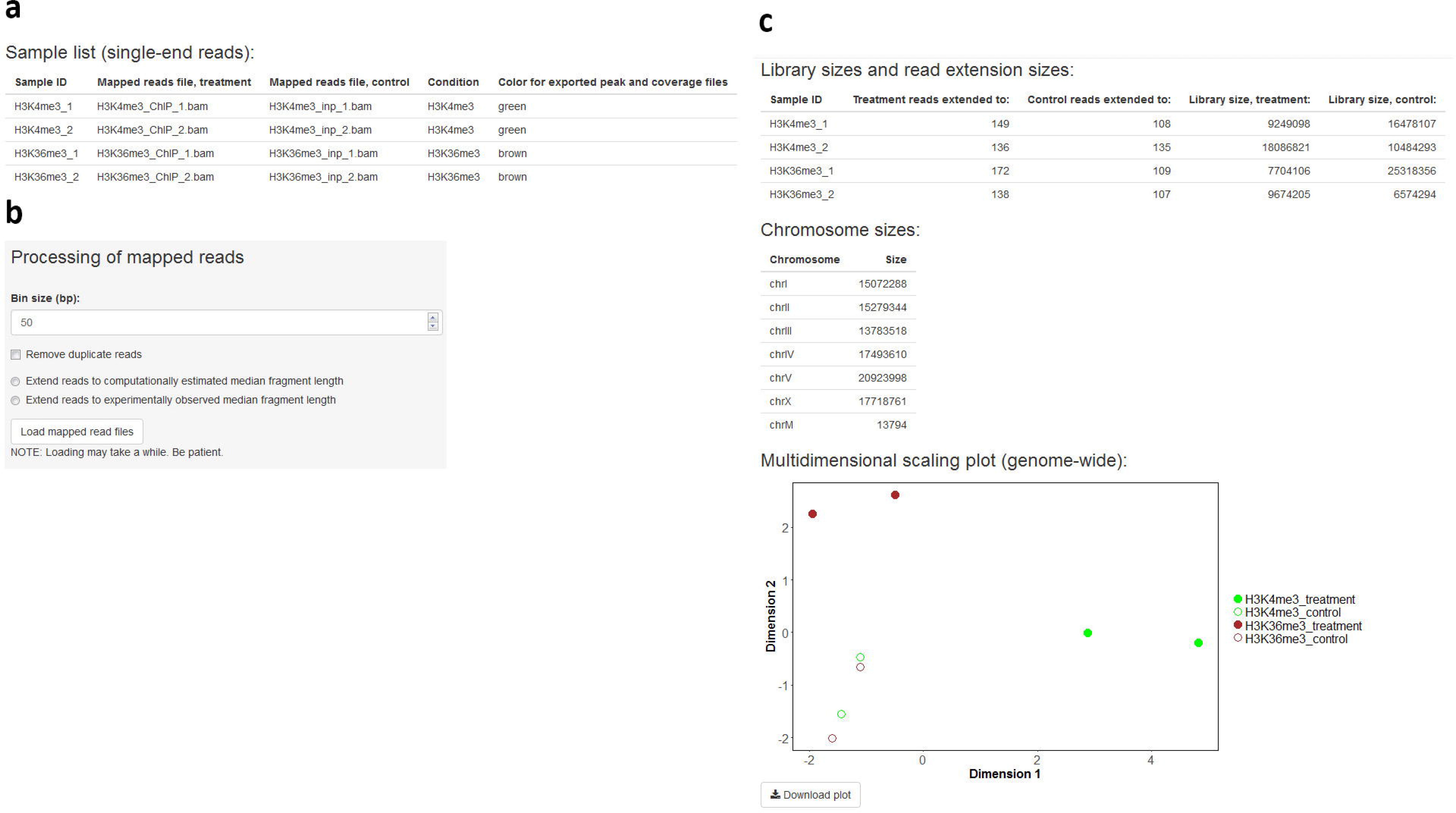
Initial processing of mapped reads. The user supplies a tab-delimited text file listing the input DNA and ChIP sample mapped read files **(a)**. If desired, duplicate read removal and fragment size extension can be performed **(b).** The application outputs a table with library sizes and median fragment sizes, a chromosome size list, and a multidimensional scaling plot representing the similarity of samples in the data set **(c)**.

Completion of the initial processing of mapped reads releases options for downstream processing which are posted to the sidebar panel, namely export of files representing sequencing coverage, peak calling and differential enrichment analysis (Fig. 4). Coverage is expressed in log2-transformed counts per million (cpm) for each genomic bin and, for each sample ID indicated by the user, three files with bedGraph format are generated: treatment (ChIP sample), control (input DNA sample) and the treatment-to-control ratio. Each file can be loaded onto a genome browser for visualization (Fig. 5). Peak calling is performed via the BayesPeak package by using a hidden Markov model and Bayesian statistical methodology [17]. The user specifies a posterior probability (PP) threshold and genomic regions with PP above such threshold are identified as peaks. If replicates are indicated, commonly enriched genomic regions representing replicated peaks can be derived, and a track definition line can be added, allowing each file to be loaded onto the UCSC genome browser for visualization. In addition, a consensus peak set representing all candidate enriched regions across the full data set may be obtained. The user may be interested in choosing this option for posterior differential enrichment analysis. Peak calling was performed for the H3K4me3 and H3K36me3 ChIP-seq data sets, yielding 4994 replicated peaks for H3K4me3 (Additional file 3) and 25855 replicated peaks for H3K36me3 (Additional file 4). Differential enrichment analysis resorts to functions implemented in the csaw [12] and edgeR [13] packages. Following library size and TMM normalization, bin-level coverage is computed and, if desired, bins in which coverage for input DNA samples exceeds that of the corresponding ChIP samples are filtered off. If the user is interested in differential enrichment analysis in a specific set of regions, the corresponding file in BED format has to be provided. To illustrate this feature of ChIPdig, differential enrichment analysis was performed for comparing H3K4me3 coverage with that of H3K36me3 using either H3K4me3 replicated peak coordinates (Fig. 6a and Additional file 5) or those corresponding to H3K36me3 peaks (Fig. 6b and Additional file 6), with a false discovery rate (FDR) threshold of 0.1. As expected, at H3K4me3 genomic peak coordinates, H3K4me3 coverage is greater than that of H3K36me3, and the opposite is observed for H3K36me3 peaks.

**Fig. 4.**
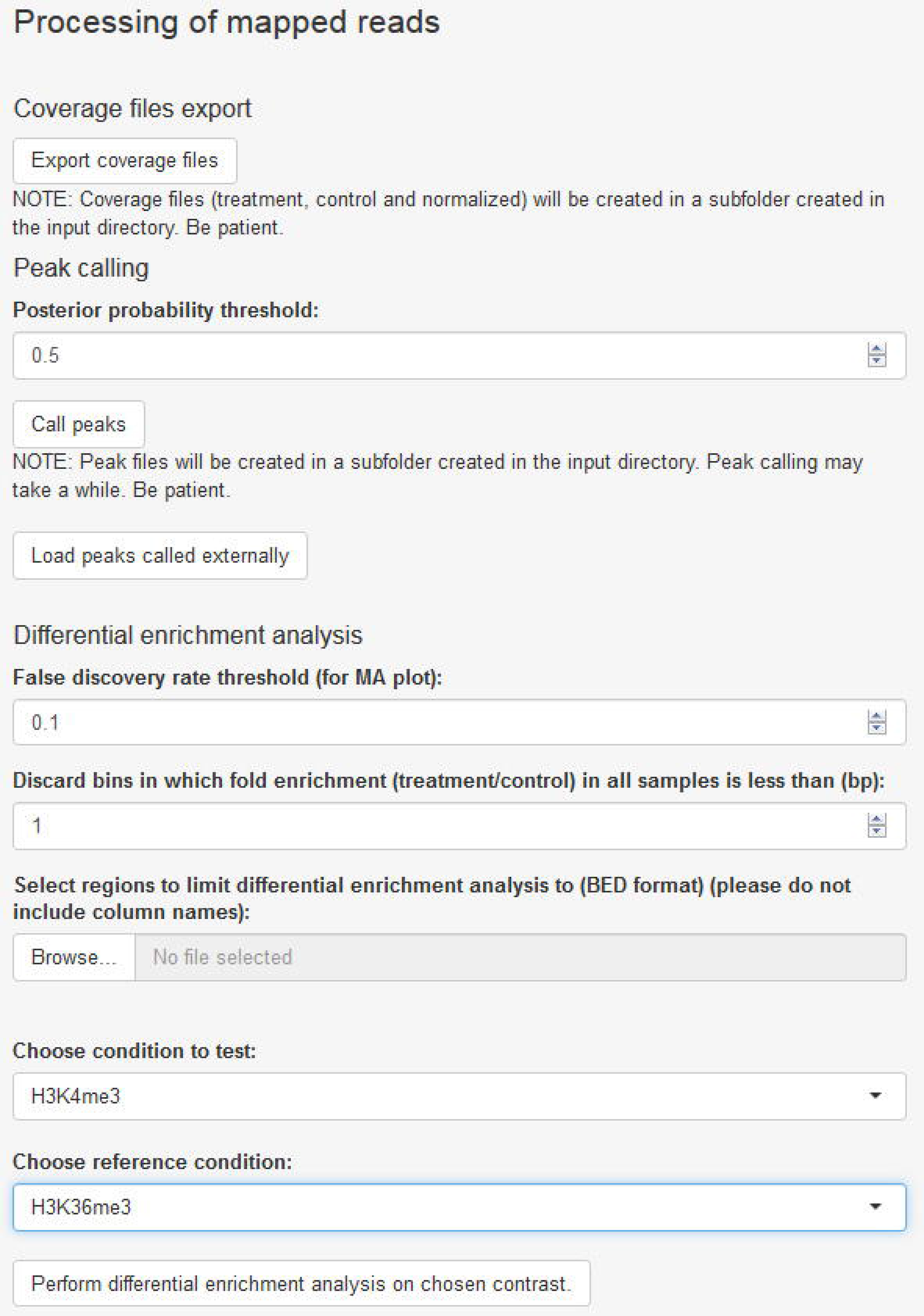
Downstream processing of mapped reads. The user may generate coverage files in bedGraph format, perform peak calling for each sample, and assess differential enrichment for a set of genomic regions of interest.

**Fig. 5.**
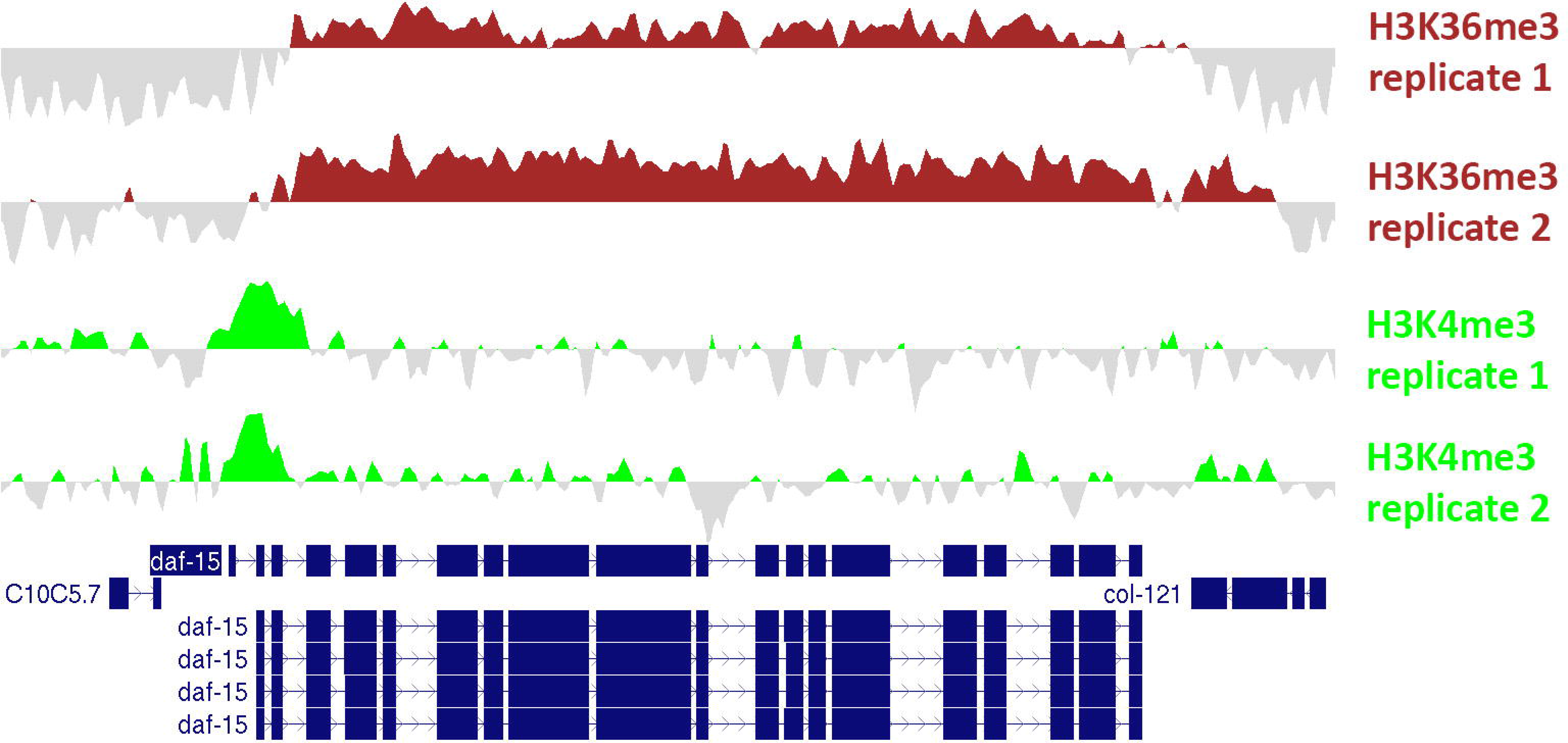
H3K4me3 and H3K36me3 ChIP-seq coverage. Values represent log2-transformed cpm for the ChIP sample subtracted by log2-transformed cpm for the input sample. Files in bedGraph format were exported from ChIPdig and loaded onto the UCSC Genome Browser. These profiles exemplify the well-known scenario whereby H3K4me3 accumulates around the transcription start site (TSS) and H3K36me3 is enriched at the gene body.

**Fig. 6.**
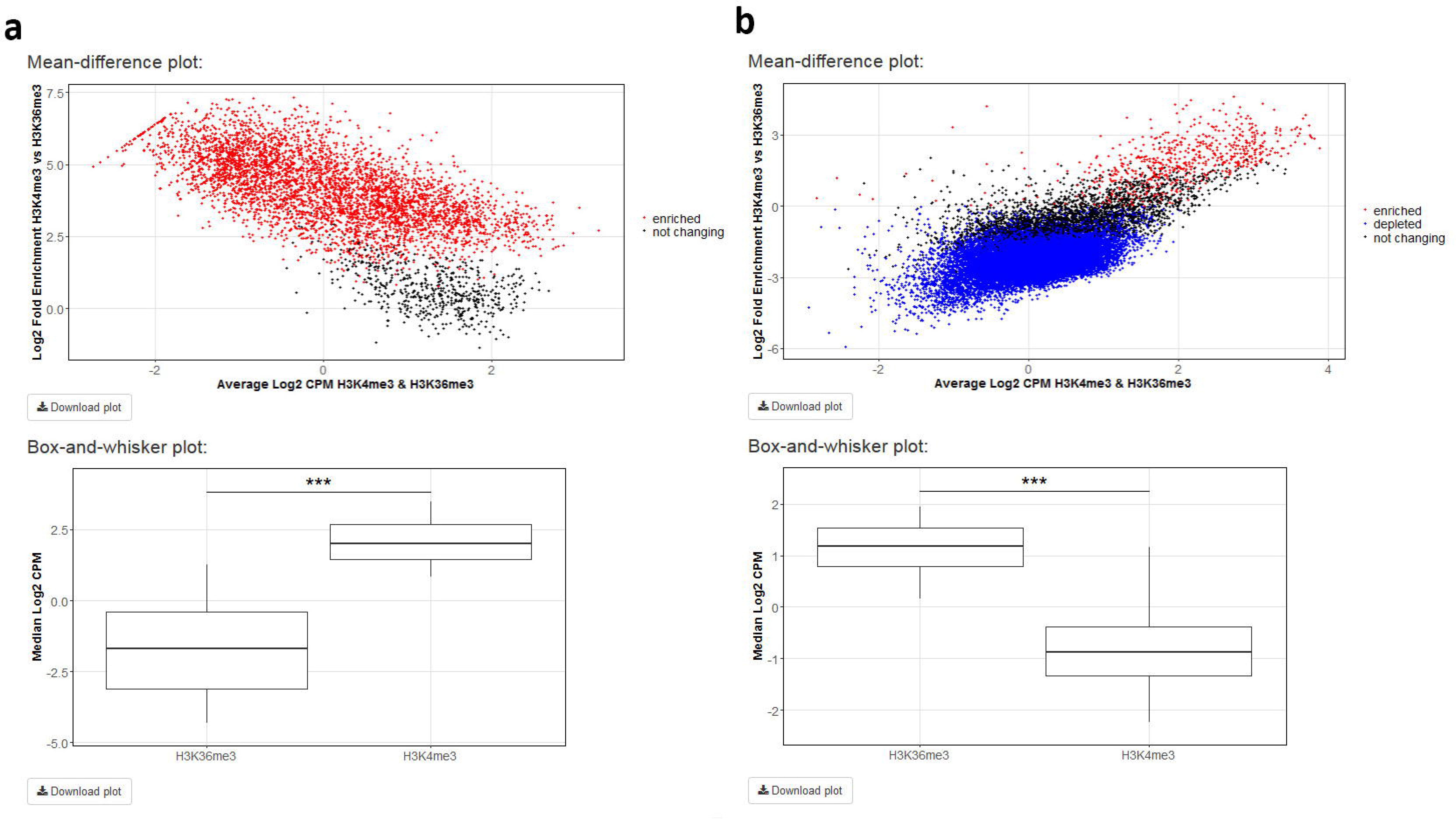
Differential enrichment analysis results. H3K4me3 ChIP-seq coverage was compared with that of H3K36me3 using either H3K4me3 peak genomic coordinates **(a)** or those of H3K36me3 peaks **(b)** with a false discovery rate (FDR) threshold of 0.1. For each analysis, a mean-different plot representing the library size-adjusted log2-transformed fold change (the difference) against the average log2-transformed coverage (the mean), as well as a box-and-whisker plot showing the global change in normalized coverage between the two conditions, were originated.

### • Annotation of genomic regions

Annotation of genomic regions of interest is performed via the ChIPseeker package [18]. The user supplies the file with regions in BED format and chooses the reference genome assembly, as well as the distance upstream and downstream of the annotated transcription start site to be considered for assignment of promoter regions (Fig. 7a). Replicated H3K4me3 and H3K36me3 peaks were annotated and, characteristically of these marks [22], most H3K4me3 peaks were assigned to promoters (Fig. 7b and Additional file 7), whereas H3K36me3 peaks lie predominantly at gene bodies (Fig. 7c and Additional file 8).

**Fig. 7.**
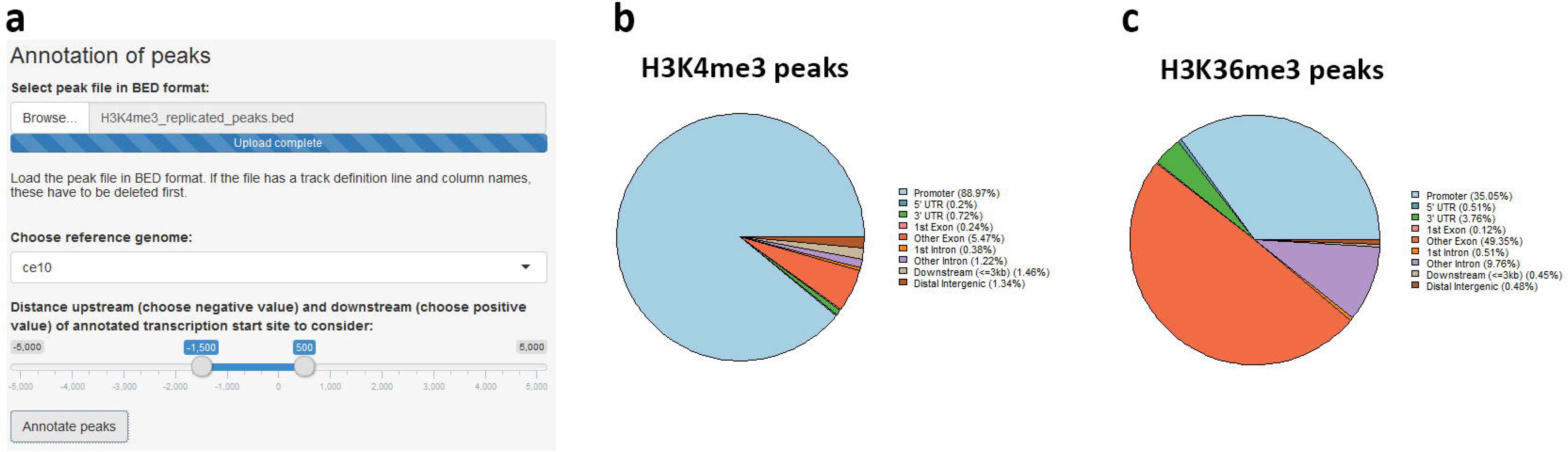
Annotation of genomic regions. The region file in BED format is uploaded and reference genome assembly, as well as distances upstream and downstream of transcription start site, are chosen **(a)**. Replicated H3K4me3 peaks are mostly promoter-proximal **(b),** whereas H3K36me3 peaks are found predominantly at gene bodies **(c)**.

### • Comparative heatmaps and metaplots

The user can supply multiple coverage files in bedGraph format and generate heatmaps and metaplots to visualize coverage in a specific region set in a comparative manner. This module of ChIPdig relies on a custom algorithm which builds a coverage matrix based on bin size and reference coordinates (start, end or both) selected by the user. Such matrix is then plotted in the form of a comparative metaplot and a set of heatmaps. H3K4me3 and H3K36me3 coverage files were loaded onto ChIPdig, along with a BED file with *C. elegans* transcription units in the WS220/ce10 reference assembly. Coverage was computed in 25 bp bins and gene bodies were either expanded or compressed to 500 bp. Windows of 250 bp upstream and downstream of each region were selected (Fig. 8a). Typical of H3K4me3, coverage is higher in the vicinity of the transcription start site, whereas H3K36me3 is enriched at transcription unit bodies.

**Fig. 8.**
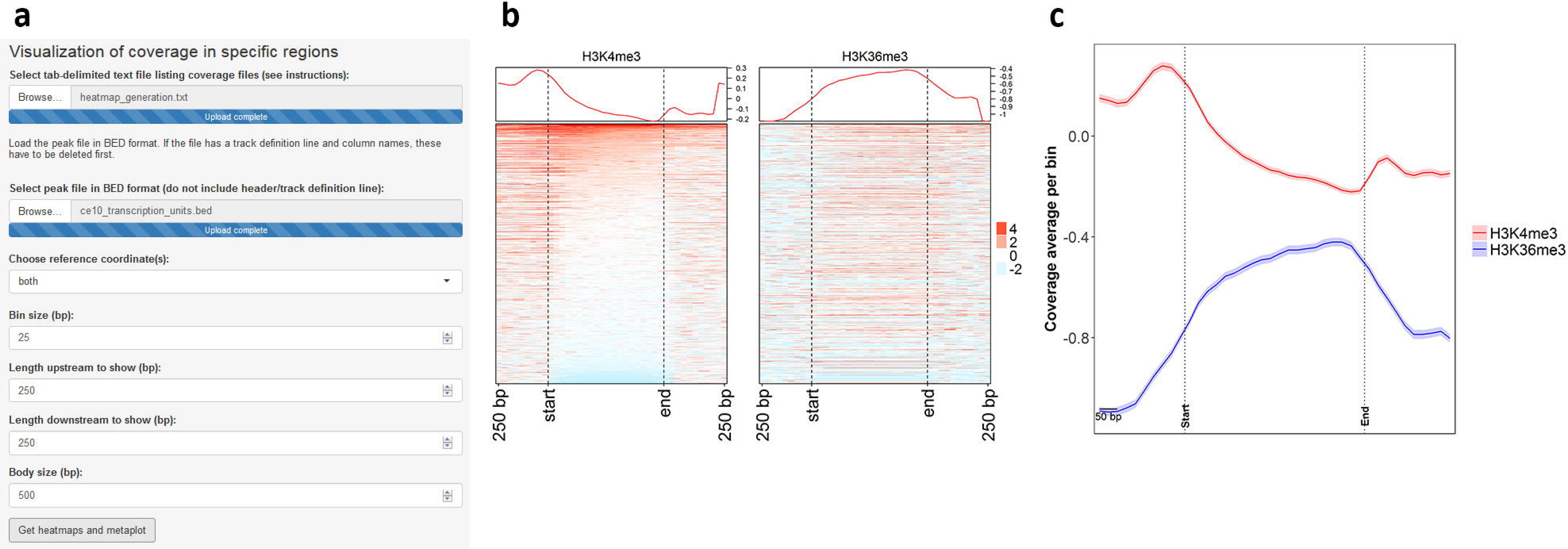
Generation of heatmaps and comparative metaplot for visualizing coverage. Coverage files for H3K4me3 and H3K36me3 were supplied to ChIPdig, as well as a file with coordinates of *C. elegans* transcription
units. The heatmaps and metaplot were generated by considering a 25-bp bin size, and gene bodies were resized to 500 bp. Windows of 250 bp upstream and downstream of each region were selected **(a).** As observed in both the heatmaps **(b)** and the comparative metaplot **(c),** coverage for H3K4me3 is overall higher in the vicinity of transcription start sites, while that of H3K36me3 is enriched at transcription unit bodies.

## Conclusions

ChIPdig is a user-friendly application for handling multiple ChIP-seq data sets and has diverse useful capabilities spanning a comprehensive analysis pipeline, namely: read alignment to a reference genome, ChIP-seq data normalization, peak calling, differential enrichment analysis, annotation of genomic regions, and generation of comparative heatmaps and metaplots for visualization of normalized coverage.

## Availability and requirements

- **Project name:** ChIPdig
- **Project home page:** https://github.com/rmesse/ChIPdig
- **Operating system(s):** Platform-independent
- **Programming language:** R, Shiny
- **Other requirements:** None
- **License:** GNU General Public License version 3.0 (GPL-3.0)
- **Any restrictions to use by non-academics:** No

## Abbreviations

bp: base pairs
ChIP-seq: chromatin immunoprecipitation followed by next-generation sequencing
cpm: counts per million
FDR: false discovery rate
GEO: Gene Expression Omnibus
GUI: graphical user interface
H3K36me3: histone H3 trimethylated at lysine 36
H3K4me3: histone H3 trimethylated at lysine 4
NCBI: National Center for Biotechnology Information
NGS: next-generation sequencing
PCR: polymerase chain reaction
PP: posterior probability
TMM: trimmed means of M values
TSS: transcription start site
UCSC: University of California Santa Cruz

## Acknowledgements

Not applicable.

## Availability of data and materials

All source code has been made publicly at https://github.com/rmesse/ChIPdig.

## Authors’ contributions

RE conceived the study and wrote the code. RE and AG wrote the paper. All authors read and approved the final manuscript.

## Consent for publication

Not applicable.

## Ethics approval and consent to participate

This study does not involve humans, human data or animals.

## Additional file description

**Additional file 1** File name and extension: Additional_file_1_alignment_report.pdf Title of data: Read alignment report Description of data: Report generated by the read alignment module of ChIPdig

**Additional file 2** File name and extension: Additional_file_2_available_colors.txt Title of data: Available colors Description of data: List of R built-in colors available for the user to choose from

**Additional file 3** File name and extension: Additional_file_3_H3K4me3_replicated_peaks.bed Title of data: H3K4me3 replicated peaks Description of data: Peaks found in both H3K4me3 ChIP-seq experimental replicates

**Additional file 4** File name and extension: Additional_file_4_H3K36me3_replicated_peaks.bed Title of data: H3K4me4 replicated peaks Description of data: Peaks found in both H3K36me3 ChIP-seq experimental replicates

**Additional file 5** File name and extension: Additional_file_5_H3K4me3_vs_H3K36me3_at_H3K4me3_peaks.csv Title of data: Differential enrichment analysis of H3K4me3 versus H3K36me3 at H3K4me3 peaks Description of data: Table summarizing the results of the differential enrichment analysis comparing H3K4me3 coverage with H3K36me3 coverage at replicated H3K4me3 peaks

**Additional file 6** File name and extension: Additional_file_5_H3K4me3_vs_H3K36me3_at_H3K36me3_peaks.csv Title of data: Differential enrichment analysis of H3K4me3 versus H3K36me3 at H3K36me3 peaks Description of data: Table summarizing the results of the differential enrichment analysis comparing H3K4me3 coverage with H3K36me3 coverage at replicated H3K36me3 peaks

**Additional file 7** File name and extension: Additional_file_7_H3K4me3_replicated_peaks_annotation_table.csv Title of data: Annotation of H3K4me3 peaks Description of data: Annotation table for H3K4me3 replicated peaks considering 1500 bp upstream and 500 bp downstream of transcription start site for promoter assignment and using *C. elegans* reference assembly ce10

**Additional file 8** File name and extension: Additional_file_7_H3K36me3_replicated_peaks_annotation_table.csv Title of data: Annotation of H3K36me3 peaks Description of data: Annotation table for H3K36me3 replicated peaks considering 1500 bp upstream and 500 bp downstream of transcription start site for promoter assignment and using *C. elegans* reference assembly ce10

